# Spatial patterns in phage-*Rhizobium* coevolutionary interactions across regions of common bean domestication

**DOI:** 10.1101/2020.07.21.214783

**Authors:** Jannick Van Cauwenberghe, Rosa I. Santamaría, Patricia Bustos, Soledad Juárez, Maria Antonella Ducci, Trinidad Figueroa Fleming, Angela Virginia Etcheverry, Víctor González

## Abstract

Bacteriophages play significant roles in the composition, diversity, and evolution of bacterial communities. Despite their importance, it remains unclear how phage diversity and phage-host interactions are spatially structured. Local adaptation may play a key role. Nitrogen-fixing symbiotic bacteria, known as rhizobia, have been shown to locally adapt to domesticated common bean at its Mesoamerican and Andean sites of origin. This may affect phage-rhizobium interactions. However, knowledge about the diversity and coevolution of phages with their respective *Rhizobium* populations is lacking. Here, through the study of four phage-*Rhizobium* communities in Mexico and Argentina, we show that both phage and host diversity is spatially structured. Cross-infection experiments demonstrated that phage infection rates were higher overall in sympatric rhizobia than in allopatric rhizobia except for one Argentinean community, indicating phage local adaptation and host maladaptation. Phage-host interactions were shaped by the genetic identity and geographic origin of both the phage and the host. The phages ranged from specialists to generalists, revealing a nested network of interactions. Our results suggest a key role of local adaptation to resident host bacterial communities in shaping the phage genetic and phenotypic composition, following a similar spatial pattern of diversity and coevolution to that in the host.

## Introduction

Bacterial viruses or bacteriophages are the most diverse and abundant biological entities on earth [1, 2]. They play a significant role in bacterial ecology and evolution, enabling horizontal gene transfer, and influencing bacterial diversity through their lytic or lysogenic cycles [3–6]. Nevertheless, phage biogeography, including patterns of dispersal, establishment, and community assembly, is very poorly understood [7], and its study could contribute to the advancement of microbiome engineering in agricultural and medical settings.

Most of the available knowledge about phage genetic diversity and spatiotemporal dynamics comes from metagenomic studies on marine cyanophages and gut microbiomes [8–10]. These studies have shown both cosmopolitan [1, 11–13] and habitat-specific phage lineages [14–21]. Given the predatory nature of phages, the presence or absence of suitable bacterial hosts shapes their distribution [8, 22–24].

Bacteriophages tend to be locally adapted to sympatric bacteria [25–28]. It is commonly predicted that local adaptation is a significant underlying factor of compositional differences across phage communities [16, 24, 26, 27, 29]. Local adaptation is a process that results in a local population of a given species exhibiting higher fitness in its local environment than in allopatric populations and takes place when environmentally driven selection is more robust than migration [30, 31]. The implied lower fitness experienced by immigrants may limit gene flow and increase genetic differentiation across populations (i.e., “isolation-by-adaptation,” or more general “isolation-by-environment”) [32–35]. Host bacteria are a key environmental driver of phage local adaptation, or more precisely, “local coadaptation”, as phage adaptation (*e.g*., increased host range) readily invokes the counter- or coadaptation of bacteria (*e.g.,* increased resistance) [25–27, 36–38].

While bacteriophages coadapt with their bacterial hosts, symbiotic bacteria coadapt with their eukaryotic hosts [39–41]. For instance, it is well established that rhizobia coadapt with legumes and form a mutualistic relationship in which they provide legumes with a steady supply of plant-usable nitrogen via nitrogen fixation [42, 43]. Aguilar et al. [44] found that Mesoamerican and Andean common bean (*Phaseolus vulgaris*) genotypes were preferentially associated with Mesoamerican and Andean rhizobia, respectively, indicating local coadaptation [44] across the two domesticated common bean gene pools [45, 46]. Furthermore, the *Rhizobium* communities were genetically differentiated, as the relative abundance of different types of the symbiotic *nodC* gene varied across the two bean gene pools [44]. Additionally, in other legume-rhizobium systems, host legume population differentiation has led to rhizobium population differentiation [47, 48] and even local adaptation [49, 50].

Symbiotic organisms have significant reciprocal evolutionary effects on each other, which in turn affect third-party interactions [51–59]. Bacterial adaptation to plant hosts may affect bacteria-phage interactions (*e.g*., by altering the expression of surface receptors for plant interactions, which serve as anchor points of phages [60–62]). Here, we predicted that the genetic differentiation and local adaptation experienced by rhizobia across common bean gene pools shape the interaction and genetic differentiation of communities of phages infecting common bean-nodulating rhizobia. Even under the assumption of high phage dispersal capabilities, we expected communities of phages to be locally adapted, showing higher infection rates for sympatric rhizobia than for allopatric rhizobia. Hence, we aimed to elucidate whether the genomic identities and host ranges of phages infecting common bean-nodulating rhizobia are geographically structured. Our approach was to sample *Rhizobium* strains and associated bacteriophages from two common bean fields in Mexico and Argentina, corresponding to independent areas of common bean domestication [45, 46]. Host species identity and bacteriophage genomic types were determined by Sanger sequencing and by both Sanger sequencing and whole-genome sequencing, respectively. Host range assessment results were analyzed for biogeographic signals and local adaptation. More specifically, we aimed to (i) assess bacteriophage diversity associated with common bean-nodulating rhizobia; (ii) compare the bacteriophage community composition across common bean gene pools; (iii) determine how bacteriophage host range pertains to bacteriophage provenance and the host species community composition; and (iv) obtain evidence of local adaptation of bacteriophages.

## Materials and methods

### Soil and *Rhizobium* sampling

The sites of rhizobia and phage sampling were common bean (*Phaseolus vulgaris*) agriculture fields located in Mexico (Tepoztlán and Yautepec, Morelos) and Argentina (Chicoana and Salta, Salta) (Fig. S1a). The bean fields were named after the municipalities in which they were located except for the ‘Salta’ field, which was located in Cerrillos near the border with the city of Salta. The distance between Tepoztlán and Yautepec is c. 7.4 km; the distance between Salta and Chicoana is c. 20.8 km. In each bean field, we collected >1 L of rhizosphere soil from three plants separated by 10 m. Each soil sample was mixed and split into two aliquots, one used for phage isolation and one used to “trap” rhizobia in the root nodules of bean plants. In Tepoztlán, three bean plants (*P. vulgaris* var. Negro Veracruz) were collected directly from the field, and their nodules were used to isolate rhizobia. To trap rhizobia from Yautepec, we used *P. vulgaris* var. Negro Veracruz, while rhizobia from Argentina were caught using *P. vulgaris* var. Alubia Cerrillos. These cultivars belong to the Mesoamerican and Andean gene pools of domesticated common bean, respectively. Previously, axenically germinated seedlings were planted in presterilized vermiculite pots and inoculated with a 100 mL soil sample. Nodules were harvested after eight to twelve weeks of growth in a greenhouse under natural environmental conditions regarding light and temperature (Supplementary Method S1). Rhizobium strains were obtained from surface-sterilized nodules via the squashing method and streaked on PY_Nal_ agar plates (Supplementary method S2; [63]). Three subculture steps were used to purify all isolated rhizobial strains. The isolates were stored at −80°C in 50% glycerol for long-term storage.

### Isolation and purification of bacteriophages

Bacteriophages were isolated from soil samples via the enrichment protocol described previously [64] using both *Rhizobium* isolates from local sites (local collection, or LC) and *Rhizobium* from the laboratory collection (standard collection, or SC; Table S1). Briefly, dry sieved soil was suspended in PY-Nal medium at a ratio of 1:2 and incubated overnight at 30°C with shaking (250 rpm). Subsequently, the soil solution was centrifuged (10,000 g, 10 min, 4°C) to remove large particles, and chloroform was added to eliminate the remaining bacteria. Chloroform was removed by centrifugation, and the solution was filtered through a 0.22 μm Millipore filter. The soil filtrate was used to inoculate one pair-member of each *Rhizobium* strain. The other pair-member served as an uninoculated control. LC rhizobia were inoculated with the filtrate from the soil of origin, while SC rhizobia were inoculated with soil filtrate that was pooled from each of the three soil samples per bean field. Cultures of 1 ml in PY_Nal_ were incubated at 30°C (250 rpm) to an optical density of 0.2 at 620 nm (OD_620_; Beckman DU650 spectrophotometer) in 96-deep-well plates. After 20 hours of incubation, the cultures were centrifuged, and the supernatant was used to reinoculate new cultures of the same pairs of rhizobial strains. This enrichment process was repeated five times; each time, the OD_620_ value was compared between the control and the inoculated pair member. A decrease in the OD_620_ was indicative of cell lysis. The presence of phages was confirmed by plaquing a dilution series of the filtrate mixed with 2.5 mL of molten soft PY (0.65% agar) on lawns of rhizobia over solid PY_Nal_ plates, as in Carlson (2005)[65]. Lytic plaques were picked and inoculated again in the respective bacterial cultures. Two additional dilution series were performed to ensure the purity of the bacteriophage stocks. Finally, the phage dilution was treated with chloroform to remove the remaining bacteria and stored at 4°C.

### Phylogeny and taxonomic *Rhizobium* species identification

Two chromosomal genes, *dnaB* and *recA*, were partially amplified by colony PCR. The primers and PCR protocols used are described in Table S2. The PCR products were purified using the Exo-SAP cleanup protocol [66]. The purified products were sent to Macrogen for sequencing (Macrogen Inc, Seoul, Korea). Sanger sequences were edited and assembled in Genius Pro v. 6.1.2. Multiple alignments were performed using MUSCLE [67], followed by manual correction to remove ambiguously aligned regions. Phylogenetic trees were reconstructed and edited with MEGA 7 using the maximum likelihood (ML) method based on the GTR+G+ I model and 1000 bootstrap replicates. Rhizobia were clustered into sequence types (STs) based on 100% sequence identity of *dnaB* and *recA* sequences.

### Genome sequencing

We obtained the genome sequences of 100 phages from the collection chosen according to their host range differences. Phage genomic DNA was purified from phage stocks propagated in the corresponding host *Rhizobium* strains. Host DNA and RNA were eliminated using DNase and RNase. Subsequently, phages were precipitated using a PEG-8000/NaCl solution. After centrifugation (10,000 g, 20 min, 4°C), the pellet was suspended in Tris-EDTA buffer. Proteins were hydrolyzed using 4% SDS and proteinase K and precipitated by adding 3 M potassium acetate. Phage genomic DNA was precipitated with 100% isopropanol and washed with 70% ethanol twice.

Phage genome sequencing was performed with Illumina technology in a Nextgen 500 system (Unidad Universitaria de Secuenciación Masiva de DNA (UUSMD-UNAM). Genomes were assembled from trimmed [68] sequence reads using the Spades v. 3.13.1 [69], Velvet v. 1.2.10 [70] and Phred/Phrap/Consed v. 23.0 [71] software packages.

The remaining 96 phages were identified via PCR and Sanger sequencing of phage marker genes used to distinguish between phage genomic types (PGTs). These genes were identified using BPGA software to obtain common genes within the PGTs and to check their presence in the remaining PGTs [72]. The extracted sequences were used to design primers with Primer3 [73]. The selected genes, primers, and PCR protocol are described in Table S2.

### Comparative genomics

The average nucleotide identity (ANI) of all pairs of phage genomes was calculated with pyani v.0.2.9 using the ANIm MuMmer method [74, 75]. PGTs were clustered based on 80% nucleotide identity and 60% coverage of the smaller genome. The assignation of PGTs to the phage morphological families of tailed bacterial viruses (Siphoviridae, Myoviridae, and Podoviridae) was performed using the VirFam server (http://biodev.cea.fr/virfam/) [76]. PGTs assigned to Microviridae were recognized by their short genome length and through BLASTn searches against the virus database of the NCBI (https://www.ncbi.nlm.nih.gov/genome/viruses/).

### *Rhizobium* susceptibility and host range assessment

The infectivity of all isolated phages was evaluated by the spot test procedure [77] in the rhizobia isolated from all four bean fields. Although this method could overestimate the lytic properties of phages and underestimate infections due to low phage titers or different adsorption efficiencies [77, 78], the spotting of phages on lawns is an affordable method for testing a large number of pairwise interactions [79]. *Rhizobium* ‘lawns’ in double-agar-layer plates were spotted with 5 μl of each bacteriophage solution prepared from the respective phage *Rhizobium* lysates. After overnight incubation at 30°C, the plates were assessed for lysis. At least three replicates of each *Rhizobium*-phage combination were performed to ensure reproducibility, and at least two replicates with the same lytic or resistant phenotype were considered to indicate a positive or negative result. Spots resulting from lysis with a translucent appearance rather than a transparent appearance were recorded as ‘partial lysis’ but were treated equivalently to transparent spots for the statistical and BiMat analyses. The binary interaction matrix is available from Github (github.com/jvancau/interactiondata).

With this information, a matrix was constructed based on Bray-Curtis dissimilarity calculated with the *vegdist* function of the *vegan* package in R [80]. To compare the phenotypic composition with the genetic composition, we clustered the bacterial hosts showing >80% similarity in terms of their susceptibility to phages into *Rhizobium* phenotype groups (RPGs, Table S3). Then, phages that infected a common range of hosts (Bray-Curtis similarity > 80%) formed phage phenotype groups (PPGs, Table S3).

### Network structure of phage-bacterium interactions

To analyze the modularity and nestedness properties of the phage-bacterium bipartite network, we employed the BiMat program [81], which maximizes the similarities in the bacteria-phage lytic interaction matrix. The program ran in the MATLAB environment and was used according to the author’s start guide [81] (https://www.github.com/cesar7f/BiMat). Modularity was tested using the Adaptive Brim algorithm, and nestedness was tested using the NODF (Nestedness metric based on Overlap and Decreasing Fill) and NTC (Nestedness Temperature Calculator) algorithms. Statistical analysis was performed using 1000 replicates and the equiprobable null model.

### Local adaptation

*Rhizobium* susceptibility rates were calculated for each *Rhizobium* strain as the number of phages able to infect the strain divided by the total number of tested phages. Similarly, phage infection rates were calculated as the proportion of rhizobia that a given phage could infect among the total number of strains tested. These values were calculated for sympatric and allopatric phage-strain combinations. The statistical significance of the differences between sympatric and allopatric susceptibility/infection rates was tested using a generalized linear model in R with a quasi-binomial model [82].

Phage local adaptation was also calculated according to Vos et al. [27] as the mean difference in the mean of the rates of sympatric (S) and allopatric (A) phage infections.

### Statistical analyses

To test the significance of compositional differences in *Rhizobium* sequence types (STs) and phage genomic types (PGTs) across common bean fields, we performed PERMANOVA tests based on Jaccard and Bray-Curtis distances with 999 permutations using the *ecodist* and *vegan* packages [80]. Principal coordinates of neighbor matrices (PCNM), which are orthogonal spatial variables derived from a spatial distance matrix, were calculated from the geographical coordinates using the ‘*pcnm’* function of the *vegan* package for R [83]. The first PCNM value was used as a proxy for spatially related variation across the two regions and was fit on a principal coordinates analysis (PCoA) using *envfit* (‘*vegan’*) with 9999 permutations. This was approach was employed to assess the significance of the distance between the two regions in explaining the compositional differences among bean fields. For each distance matrix type, the correlations among all datasets of the genetic and phenotypic composition across bean fields (ST, PGT, RPG and PPG) were assessed using Mantel tests (999 permutations, *vegan* package).

A Mantel test was also used to correlate *Rhizobium* and phage genetic distances with the corresponding susceptibility range and host range distances across all isolated rhizobia and phages. PERMANOVAs were used to test the significance of the effects of the origin and genetic or taxonomic identity of rhizobia and phages on their susceptibility range and host range, respectively.

### Phage and *Rhizobium* nucleotide accessions

Phage genome sequences are available from GenBank with IDs MN988459 to MN988558. Rhizobium nucleotide sequences are available under: MT756388 – MT756428 (*dnaB*) and MT756429 – MT756469 (*recA*).

## Results

### *Rhizobium* and phage community sample composition

We studied 229 *Rhizobium* strains isolated from four agricultural plots in Central Mexico (Tepoztlán and Yautepec) and Northwest Argentina (Salta and Chicoana) (Fig. S1a-b). The rhizobial strains from each site (LC, local collections) were employed to trap phages from the soil of the same locality by the enrichment method [64]. In parallel, the same soils were pooled for each plot and used to search for phages using a standard collection (SC) of 94 *Rhizobium* strains of diverse geographic origins maintained in our laboratory (Table S1). A total of 196 phages were obtained with this protocol, 110 from LC and 86 from SC (Fig. S1b).

### Genetic differentiation of *Rhizobium* populations between agricultural fields

Our first aim was to investigate whether the collected *Rhizobium* strains represent geographically structured populations. Phylogenetic analysis of the partial *recA*-*dnaB* sequences of 229 rhizobial strains identified *R. etli* as the predominant species at Mexican sampling sites (74.6%), while *R. phaseoli* was dominant in Argentina (80%) (Fig. 1; Fig. S2a). According to the nucleotide variations in *recA* and *dnaB*, all of the isolated rhizobia were grouped into 41 chromosomal sequence types (STs) (Table S4). PERMANOVA tests employing two distance measures (Jaccard and Bray-Curtis) indicated that the rhizobial communities differed significantly in terms of genetic composition and the relative abundance of genotypes (STs) among different bean fields and regions (Fig. 2a-c; Table S4-S5). The geographic distance between the regions, represented by a PCNM vector, was correlated with the ST composition among the rhizobia isolated from the four common bean fields (Table S5). At the species level, *R. etli* and *R. phaseoli* STs differed significantly among the sites of origin of the common bean fields. Between regions, only the *R. etli* populations differed in terms of the ST composition, whereas the *R. phaseoli* ST composition changed only marginally across regions (Table S5). The abundance of ST5 in *R. phaseoli*, the only ST of the 41 total STs found across the four common bean fields, might explain this last result (Fig. 2c; Table S4).

**Fig. 1.**
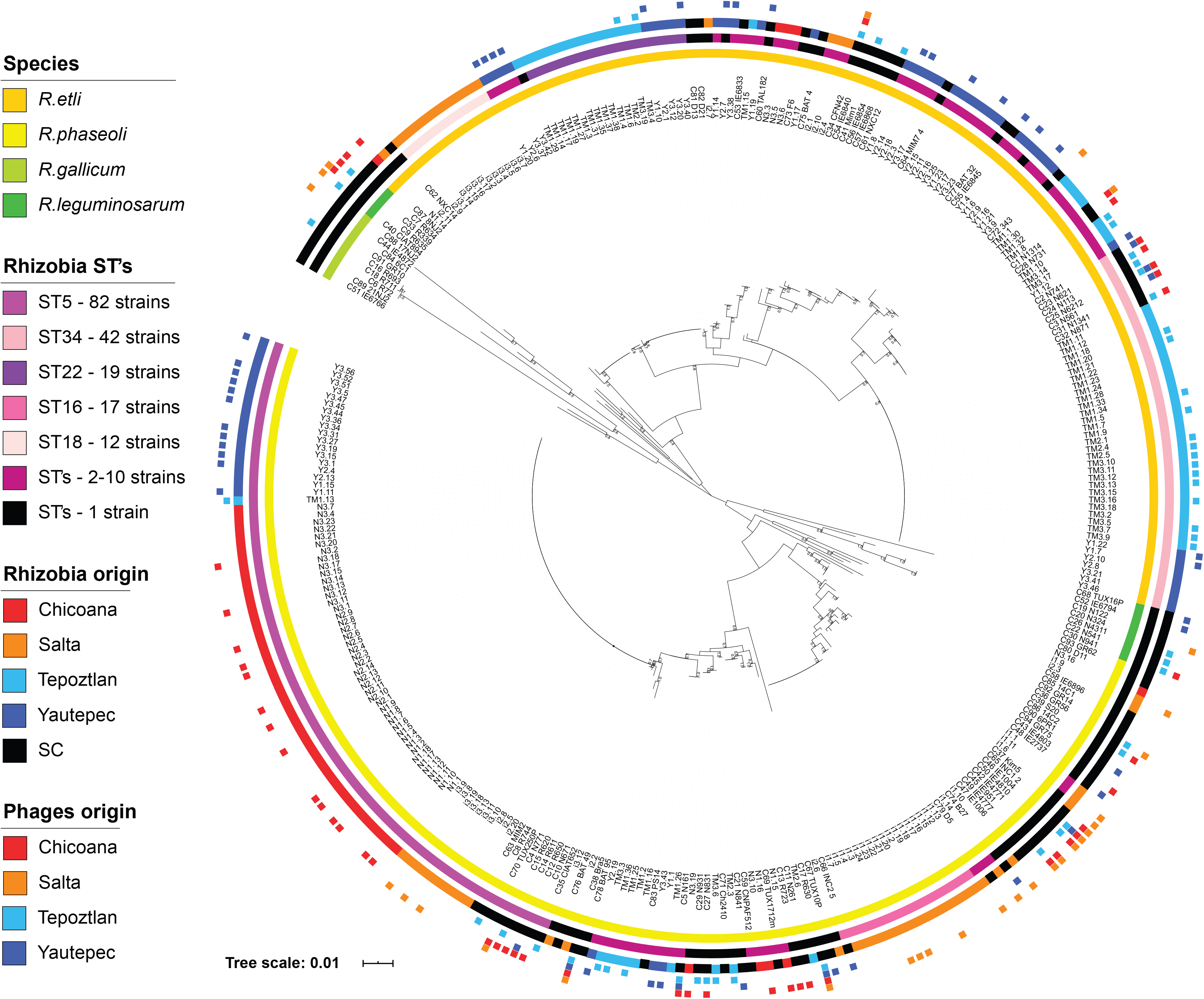
Phylogenetic tree of all rhizobia collected from Mexico and Argentina (LC, n = 229) and from the standard laboratory collection (SC, n = 94). The tree was constructed using the maximum likelihood method and is based on the concatenated sequences of two chromosomal genes (dnaA-*recA*). The tree bar scale indicates the number of nucleotide substitutions per site. Insets on the left side explain the contents of the four concentric circles. From the inner to the outermost circles: taxonomic classification of rhizobia, *Rhizobium* chromosomal STs (sequence types; no STs were assigned to SC strains), field of origin of the strains, and squares indicating the origin of the phages isolated using the corresponding strain.

**Fig. 2.**
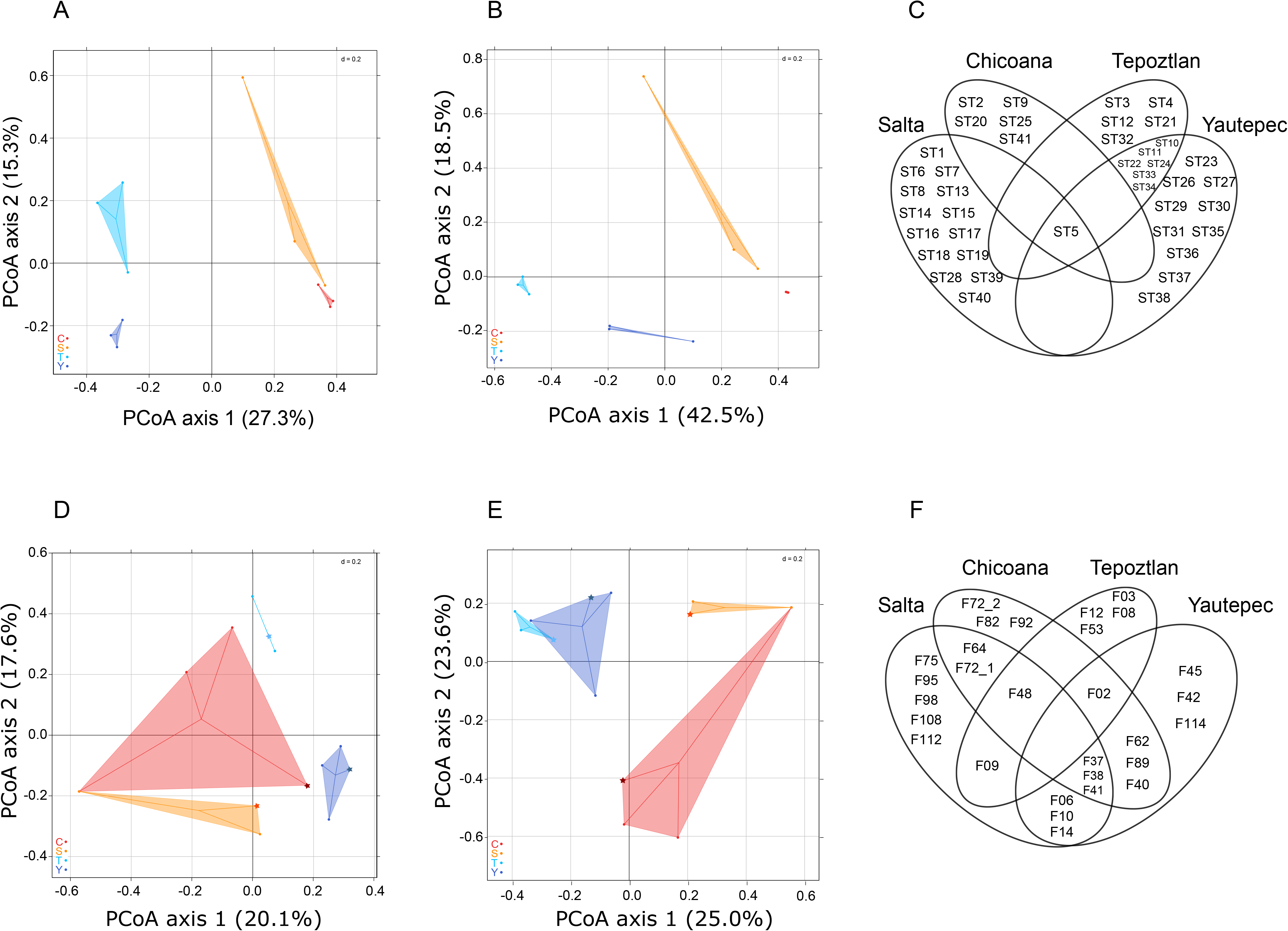
Differentiation of *Rhizobium* and phage communities. A and B, PCoA plots illustrating the differences in chromosomal composition between *Rhizobium* communities across common bean populations based on Jaccard distances (presence-absence) and Bray-Curtis distances (relative abundance), respectively. D and E, PCoA plots showing the differences in the phage genomic type composition between phage communities across common bean populations based on Jaccard distances and Bray-Curtis distances. Bean fields are indicated by different colors: Tepoztlán (T), light blue; Yautepec (Y), dark blue; Salta (S), orange; Chicoana (C), red. Phages isolated by inoculating pooled samples from a given location into rhizobia from the laboratory’s standard collection of rhizobia (SC) are indicated with a star symbol. C and F, Venn diagrams of the distribution of *Rhizobium* chromosomal STs and phage genomic types (PGTs), respectively.

### Diversity of phage communities

To assess phage diversity, we obtained the complete genome sequences of 100 out of 196 phages. The phage genomes displayed a wide range of lengths (from 4.8 to 207.6 kb; median 54.4) and GC contents (from 41% to 61%; median 57%). They were clustered into 29 PGTs (defined by ANIm, see Methods), 18 of which had two or more individual genomes, and 11 were singletons. Within PGTs, the genomes exhibited nucleotide variation ranging from 85.8 to 99.7%, with a coverage of approximately 64.1 to 100% (Fig. S3). Additionally, 90 phages among the remaining 96 phages were assigned to PGTs by the PCR identification of specific phage genes conserved within the members of the 29 PGTs, followed by Sanger sequencing (see Methods) (Table S2).

Phages were also classified into morphological families using VirFam predictions [76]. Most of them (26/29) belonged to the order Caudovirales, represented by the families Podoviridae (12 PGTs), Siphoviridae (8 PGTs), and Myoviridae (6 PGTs). Three PGTs were identified as members of the Microviridae family.

Most phage families were present at the four sampling sites, but the Salta community was dominated by Myoviridae (69%) and the Chicoana community by Siphoviridae (62%) (Fig. S2b; Table S6). Remarkably, the Microviridae family (F02), defined by small-genome phages (4.8 – 6.2 kb), was dominant in Tepoztlán (60%), whereas it showed low abundance or was absent in the other populations (0-23%; Table S6).

### Phage population differentiation across agricultural sites

Following the differences in the rhizobial ST composition per sampling site, we found that the composition of PGTs also differed significantly among common bean fields and regions (Fig. 2 d-f, Table S6). The differences were significantly correlated with geographic distance (Table S5). The phage communities were also significantly different between the Mexican bean fields within regions based on Jaccard distances (F_1,5_ = 3.744, P = 0.024), but they were not significantly different between Mexican bean fields based on Bray-Curtis distances (F_1,5_ = 3.679, P = 0.055) or between Argentinian bean fields (Jaccard: F_1,5_ = 1.132, P = 0.384; Bray-Curtis: F_1,5_ = 2.204, P = 0.149). Mantel tests showed that the genetic composition differences among phage communities were significantly correlated with the differences in the *Rhizobium* community genetic composition among bean fields (Table S5). Some PGTs coexisted at two of the sampling sites, whereas PGTs rarely coexisted at three sites and never at four (Fig. 2f). Moreover, 52% of the 29 PGTs occurred solely in one bean field, and 69% were restricted to a particular region. A spatial pattern distinction was also shown by the ANI values, as the average ANI of allopatric phages belonging to the same PGTs was 88%. In comparison, the average ANI of sympatric phages belonging to the same PGT was 96%.

### Structure of the phage-bacterium interaction network

To define the rhizobium-phage interactions within and between the four communities, we tested the infectivity of 196 phages against 229 *Rhizobium* strains by the spot assay (see methods). We registered the following phenotypes: complete lysis (transparent spots), partial lysis (translucent spots), and resistance (absence of lysis), in three independent experiments. A total of 44,884 interactions were examined in triplicate experiments; 19,474 plaques showed full or partial lytic phenotypes, recorded as positive interactions (1 in the bipartite network of Fig. 3) [81, 84]. The rhizobium-resistant phenotypes were recorded as 0 in the corresponding binary matrix (Fig. 3).

**Fig. 3.**
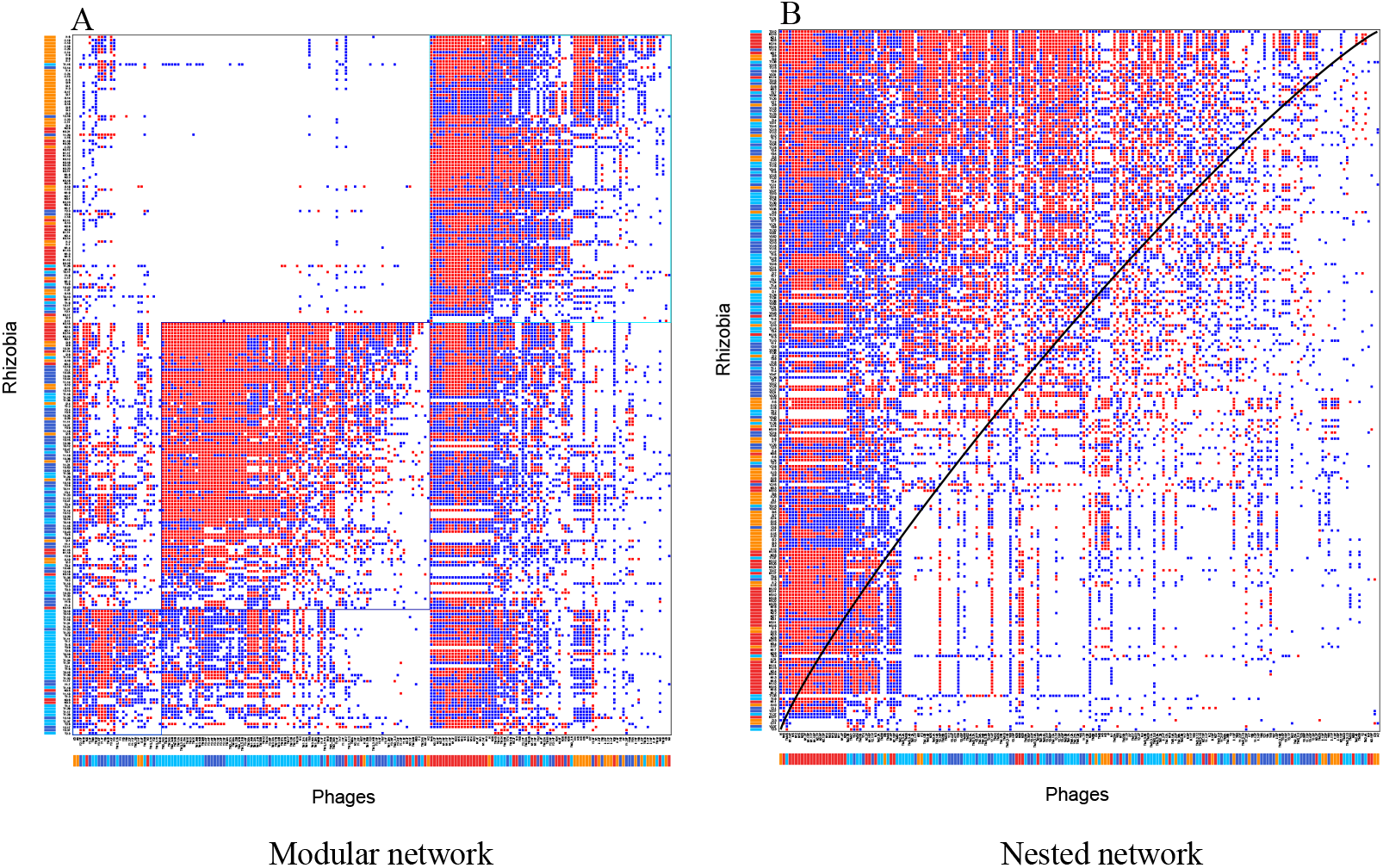
Structure of phage-*Rhizobium* interactions. Phage–bacterium infection network of interactions between 196 phages (columns) and 229 *Rhizobium* strains (rows) performed with BiMat [81]. Either full (red) or partial (blue) lytic interactions were recorded as positive interactions in at least 2/3 of independent tests; blank cells indicate the absence of interaction. In the bottom row, colored bars indicate the bean field of origin of the phages: Tepoztlán (T, Mexico) = light blue; Yautepec (Y, Mexico) = dark blue; Salta (S, Argentina) = orange; Chicoana (C, Argentina) = red. The first column on the right indicates the site of origin of rhizobia with the same colors of the bars used for the phages. (A) Modular sorting of the interaction data, visualizing the presence of three modules. (B) Nested sorting of the interaction data, visualizing the spectrum of generalists to specialist phages.

Overall, the BiMat network showed high connectance (0.43), indicating that there were approximately 4/10 effective phage-bacterium interactions in the community, and weak modular differentiation (Qb = 0.21; Fig. 3a). Three large modules, with high internal connectance in comparison with the outside modules, were detected (Fig. 3a). Phages from Mexico (Tepoztlán and Yautepec) generally interacted best with *Rhizobium* isolates from Mexico. It was less common for these phages to infect *Rhizobium* from Argentina. These results suggest large-scale modularity dominated by sympatric interactions. However, some exceptions were observed; for instance, phages from Chicoana were very promiscuous, since a significant number of Mexican strains were cross-infected by these Argentinian phages (Fig. 3a). This may in part explain the overall nested structure shown by the BiMat matrix, quantified by the independent NODF (0.68) and NTC (0.73) algorithms. This suggests that the phage communities consist of a range of specialist to generalist phages (Flores et al. 2013; Fig. 3b).

### *Rhizobium* susceptibility and phage host range

Although all *Rhizobium* strains were closely related in the *recA*-*dnaB* phylogenetic tree, they varied widely in their susceptibility range, with some rhizobia being infected by approximately 10.2% to 74.0% of the phages (average rate of infection = 43.4%). Hence, we sought to investigate whether the susceptibility of rhizobia was associated with geographic origin, taxonomic affiliation or genetic identity. When we compared the variation in the susceptibility range between individual rhizobia, we found that the susceptibility of rhizobia was significantly different not only between species (F_1,217_=10.296, P=0.001) (Fig. 4a) but also among bean fields (F_3,217_=30.964, P=0.001), and we observed the interaction of the two factors (F_3,217_=6.3575, P=0.001; Fig. 4a). The last finding indicates that the effect of *Rhizobium* species identity on the susceptibility range depends on the geographic origin of the rhizobia. When species were split into STs, the *Rhizobium* susceptibility range was found to differ significantly among bean fields (F_3,178_= 53.458, P=0.001) and STs (F_40,178_=6.056, P=0.001), although their interaction was not significant (F_6,178_=0.955, P=0.542) due to the limited geographic spread of most STs. Susceptible strains were clustered in 48 RPGs (*Rhizobium* Phenotype Groups; see methods). Twenty-nine RPGs were singletons, whereas 19 were formed by more than two strains (maximum of 40) (Table S3). Thirty-two RPGs belonged to *R. etli*, twelve RPGs corresponded to *R. phaseoli* (RPG4, 13, and 19), and the other four RPGs belonged to both species (RPG1, 2, 6, and 11). The most abundant RPGs (1 to 4) were composed of the frequently identified STs of *R. phaseoli* (ST5) and *R. etli* (ST10 and ST34). We found that the RPG composition was strongly correlated with the ST composition (Table S7). Additionally, *Rhizobium* genetic distance and susceptibility range similarity were significantly correlated (r= 0.3125, P= 0.001).

**Fig. 4.**
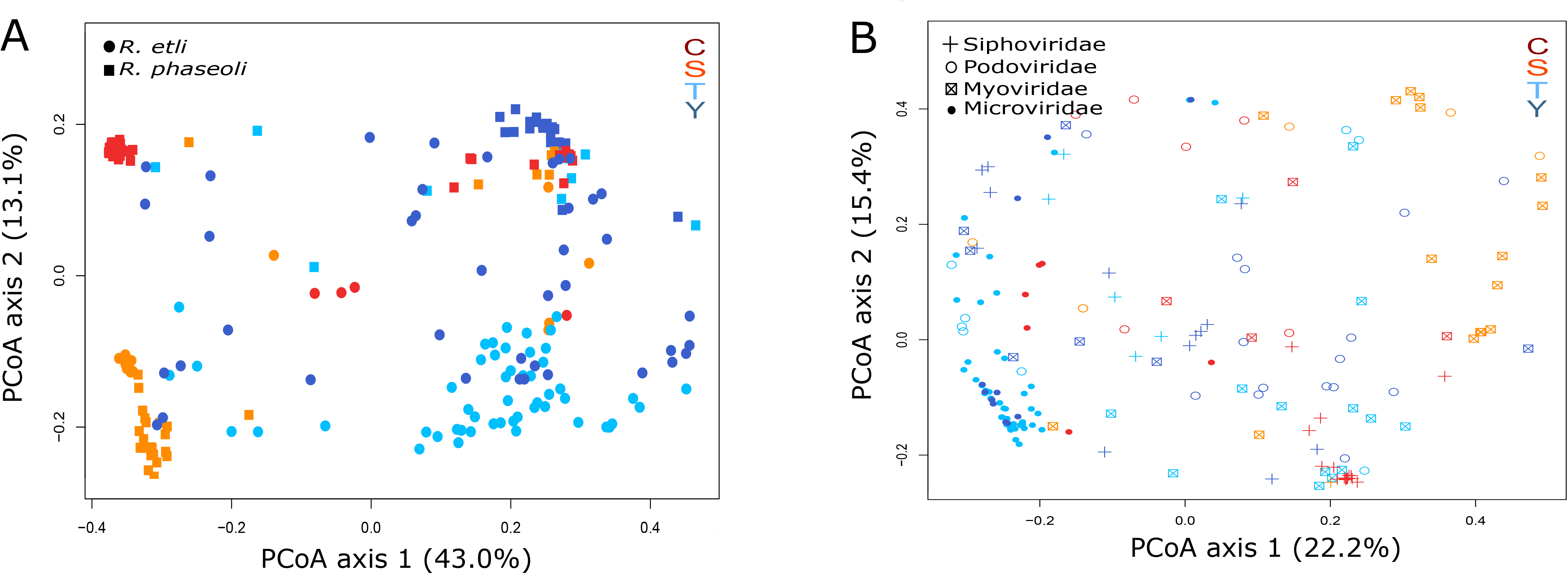
Principal coordinates analysis (PCoA) plot showing the Bray-Curtis dissimilarity in the *Rhizobium* susceptibility range among common bean fields and *Rhizobium* species in A and the Bray-Curtis dissimilarity in the phage host range among regions and phage taxonomic families in B. Each axis explains a certain fraction of dissimilarity, given within parentheses. Different *Rhizobium* species and phage taxonomic families are represented by symbols. The bean fields of origin are indicated by different colors: Tepoztlán (T), light blue; Yautepec (Y), dark blue; Salta (S), orange; Chicoana (C), red.

We examined the host range of phages to identify those with similar infection spectra. Phage infection rates, or the proportion of hosts that a given phage isolate could infect, varied considerably from 2.2% to 92.6% (average infection rate 43.4%). All but three phages were able to infect both *R. etli* and *R. phaseoli*; however, the phages were able to infect only 51% of STs on average. The host range of the phages was significantly affected by the bean field of origin (F_3,142_=22.938, P=0.001), phage genomic type (PGT; F_27,142_=6.930, P=0.001), and the interplay between these factors (F_14,142_=2.882, P=0.001). The last finding indicates that the effect of PGT on the host range depends on the geographic origin of the phage. Similar results were obtained when the phages were grouped by taxonomic family: phage host range was significantly affected by the bean field of origin (F_3,172_=14.158, P=0.001), family (F_3,172_=8.355, P=0.001), and the interplay between them (F_8,172_=3.872, P=0.001) (Fig. 4b).

Based on Bray-Curtis dissimilarity < 20%, 139 phages (70% of the total) were clustered into 24 PPGs; the remaining phages (57 phages) exhibited a unique PPG. Within the PPGs, two profiles accounted for 53% of the phages, whereas 22 profiles exhibited fewer than ten phages from similar PPGs. The PPG composition across common bean fields was significantly correlated with the RPG composition based on Bray-Curtis distances, but not based on Jaccard distances (Table S7). The PPG composition was also significantly correlated with PGT composition (Table S7). Accordingly, host range similarity was considerably correlated with phage ANI (r= 0.2890, P= 0.001).

### *Rhizobium*-phage local adaptation

The above results suggest that phage communities are adapted to *Rhizobium* in their areas of origin. To examine this issue, we estimated the infectivity rates of phage isolates and the susceptibility rates of *Rhizobium* isolates in sympatric and allopatric combinations. Despite ample variation, phage infection rates were significantly greater for sympatric infections (mean = 0.55 ± 0.01; CV = 55%; σ^2^ =k 0.092) than for allopatric infections (mean = 0.39 ± 0.02; CV = 64%; σ^2^ = 0.062; F_1,390_ = 23.234, P = 2.06e-6) (Fig. 5a), suggesting a trend of local adaptation. This was true for both phages isolated using the standard collection (SC phages; F_1,170_= 5.656, P= 0.0185) and phages isolated using local rhizobia (LC phages; F_1,218_= 20.133, P= 1.17e-05). However, sympatric phage infection rates were significantly higher than allopatric infection rates for Mexican communities (0.52 ± 0.02 *versus* 0.35 ± 0.02; F_1,252_= 30.210, P = 9.49e-08; SC: F_1,94_= 7.053, P= 9.30e-03; LC: F_1,156_= 26.405, P= 8.17e-07), but the difference was not statistically significant for Argentinean communities (0.60 ± 0.04 *versus* 0.47 ± 0.04; F_1,136_= 3.542, P = 0.062; SC: F_1,74_= 0.989, P= 0.323; LC: F_1,60_= 2.730, P= 0.104) (Fig. 5b). This was largely due to the fact that the phages isolated from Chicoana (Argentina) showed no differences between the rates of infection of local rhizobia and those of nonlocal rhizobia isolated from the other three sites (Fig. 5e). Although the Mexican phages were locally adapted overall, phage cross-infection was detected at similar rates between the communities of Tepoztlán and Yautepec (Mexico), indicating nonadaptive phage differentiation between these communities (Fig. 5e). Indeed, the infection rates of Mexican phages in Argentinian rhizobia were significantly lower, suggesting an effect of geographic distance on local adaptation (Fig. 5b).

**Fig. 5.**
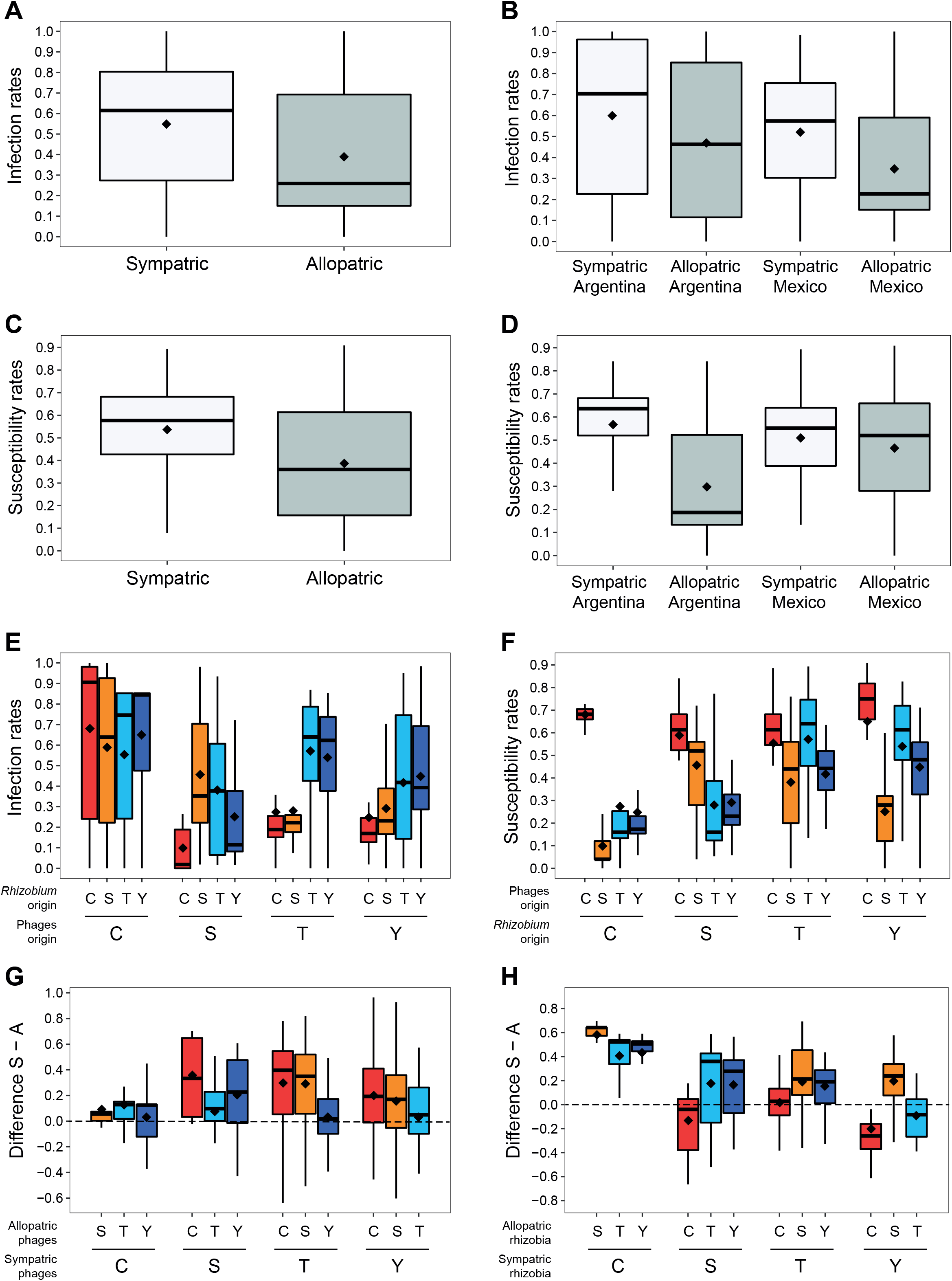
Box plots of phage infection rates and *Rhizobium* susceptibility rates indicating phage local adaptation and *Rhizobium* maladaptation. Phage infection rates between sympatric *versus* allopatric combinations averaged across all locations (A), averaged across populations within regions (B), or among populations (E). *Rhizobium* susceptibility rates between sympatric *versus* allopatric combinations averaged across all locations (C), averaged across populations within regions (D), or among populations (F). The mean is given for each box plot (black diamond). Differences between the means of sympatric (“S”) *versus* allopatric (“A”) phage infection rates (G) and the rates of the susceptibility of *Rhizobium* (H) are also shown. The origins of the phage and *Rhizobium* samples are indicated by T = Tepoztlán (Mexico), Y = Yautepec (Mexico), S = Salta (Argentina), and C = Chicoana (Argentina).

On average, the rhizobia were more susceptible to sympatric phages (mean 0.54 ± 0.01; CV= 36.9%; σ^2^ = 0.039) than to allopatric phages (mean 0.40 ± 0.01; CV= 49.7%; σ^2^ = 0.040), indicating that the bacterial populations were maladapted to the local phages (F_1,456_= 46.652, P= 2.72 e-11; Fig. 5c). Similar results were obtained regardless of whether susceptibility rates were calculated for each *Rhizobium* isolate using only infections by phages isolated by using the standard collection (SC phages; F_1,456_= 38.942, P= 9.99e-10) or phages isolated by using local rhizobia (LC phages; F_1,456_= 45.901, P= 3.85e-11). All of these results suggest that rhizobia are maladapted to coexisting phages. However, the high susceptibility of rhizobia to local phages appeared to hold only for Argentinian *Rhizobium* communities (0.57 ± 0.02 *versus* 0.30 ± 0.02; F_1,212_= 64,277, P= 7.20e-14; SC: F_1,212_= 102.51, P< 2.2e-16; LC: F_1,212_= 42.256, P= 5.64E-10), since the Mexican *Rhizobium* isolates showed no significant differences in susceptibility to sympatric and allopatric phage infection (0.51 ± 0.02 *versus* 0.49 ± 0.01; F_1,242_= 1.232, P= 0.268; SC: F_1,242_= 0.440, P= 0.508; LC: F_1,242_= 1.999, P= 0.159) (Fig. 5d; Fig. 5f). When we considered the susceptibility of *Rhizobium* to sympatric phages or allopatric phages by site, the Chicoana *Rhizobium* community was found to be very susceptible to its own phages but resistant to phages from the other sites (Fig. 5f). In contrast, the rhizobial strains from Tepoztlán, Yautepec, and Salta were very susceptible to Chicoana phages (Fig. 5f).

In most cases, the difference between the mean sympatric rates of phage infection (S) and the mean allopatric rates of infection (A) indicated phage local adaptation in the four rhizobium-phage communities (Fig. 5g). In contrast, resident rhizobia were more susceptible to sympatric than to allopatric phages, except for the rhizobia from Yautepec that were also efficiently infected by Tepoztlán phages (Fig. 5h). The four populations of rhizobia were susceptible to the Chicoana phages, but the Chicoana rhizobia were mostly resistant to allopatric phages (Fig. 5h).

## Discussion

Overall, our data indicate that the genetic and phenotypic diversity of phages and their *Rhizobium* hosts is spatially structured and that phages are adapted to their local host communities. Previous research has shown that rhizobia are spatially structured and can locally adapt to their legume hosts and other local environmental factors [49, 50, 85, 86]. For instance, across the areas of common bean domestication, rhizobia receive a greater competitive benefit when nodulating sympatric common beans (*Phaseolus vulgaris*) than nonnative common bean varieties [44]. In turn, our results show that sympatric phage communities have locally adapted to these rhizobia.

We found that each of the four phage communities analyzed was dominated by a particular taxonomic family, prominently Microviridae in Tepoztlán, Myoviridae in Salta, and Siphoviridae in Chicoana. Moreover, phage genome sequencing revealed high genetic diversity within taxonomic families. Phage genetic diversity varied considerably within and between communities, and phages were clustered into phage genomic types (PGTs). Approximately 52% of the 29 PGTs occurred solely in a single phage community. Similarly, Scola et al. [87] found that 66.4% of Namib Desert soil phage OTUs were exclusive to a single sampling site. Other PGTs (31%) occurred in both Mexican and Argentinian bean fields, with an average ANI of 88% across regions. Moreover, phages of the same PGT exhibited an 8% higher ANI within regions than between regions on average. Highly similar phages have previously been found across multiple distant aquatic ecosystems around the world, in human virome samples [88, 89] and even in more distinct ecosystems [11]. The results indicate an emerging pattern in which a higher fraction of phage community members present a limited geographic range, while a significant minority of relatively closely related phages are distributed globally [16, 87, 90]. It remains unclear to what extent such observations are due to dispersal limitation [8]. The *Rhizobium* communities also showed spatial structure. *R. etli* was the predominant species nodulating common bean in the analyzed agricultural fields in Mexico, and *R. phaseoli* predominated in the Argentinian fields, although both species were mainly characterized by spatially restricted STs.

Phage community (PGT) spatial patterns were correlated with the compositional differences among *Rhizobium* (ST) communities. Similar correlations between host-phage communities have been seen in aquatic systems [16, 17, 22, 91–93], which seems to verify the common assumption that the relative abundance of phages within a community depends largely on host abundance and susceptibility [8, 23]. However, the presence of susceptible rhizobium lineages did not imply the presence of specific phages or *vice versa*. For example, the omnipresent ST-5 was susceptible to all but two members of the phage lineages, but most phage lineages were spatially limited. Similarly, F06 phages could infect members of all STs, yet their presence was limited to Mexican bean fields. Although our study provides detailed genetic information on phages, the *Rhizobium* lineages were broadly defined by housekeeping gene markers (*recA*, *dnaB*). It is expected that much of the still-uncharacterized phenotypic and genetic microdiversity within bacterial species [94] may explain the spatial heterogeneity of the phage community composition and host-range patterns better than the presence or absence of a suitable host (defined by species or ST).

Host range breadth varied considerably among phages, from generalists to specialists, resulting in a nested structure of the inferred whole cross-interaction network. Based on the extensive analysis of experimental cross-infection data, Flores et al. [95] concluded that nestedness is the characteristic profile in most cases. A modular network structure may be significant at large phylogenetic scales [84], but genotype-to-genotype interactions are most frequent within narrow phylogenetic ranges and result from coevolutionary processes of susceptibility and resistance. High host genetic similarity may underlie the nested structure of our network.

Nevertheless, we found that phage-rhizobium interactions were significantly affected by the genetic identity of both phages and rhizobia as well as their geographic origin. This may explain the detection of three large modules with high internal connectivity and suggests ongoing local adaptation. Indeed, we showed that phages infecting common bean-nodulating rhizobia experienced higher infection rates in sympatric rhizobia than in allopatric rhizobia; hence, they were locally adapted. Furthermore, sympatric phages showed more similar host ranges than allopatric phages, and sympatric rhizobia shared similar susceptibility ranges. Only a few field studies have provided evidence that phages locally adapt to their bacterial hosts in nature [27, 84, 96]. Although the *Rhizobium* communities were generally maladapted to the local phages, they may be adapted to their local environment as a whole. A nodule may provide an isolated niche for rhizobia where they may survive competition with and the antagonistic effects of other bacteria or, more directly, phages. In free-living conditions, depredation by phages may change the population structure of rhizobia. Phages are usually considered to be slightly ahead of their host in their coevolutionary arms race due to the higher selective pressure they experience and their greater evolutionary potential [27, 97].

Although local bacterial adaptation to phages has been described multiple times in coevolutionary *in vitro* experiments [36, 98, 99], evidence in nature appears to be lacking [26]. This discrepancy is probably due to the relatively high availability of resources to hosts *in vitro*, which sways the arms race to the benefit of the host [100].

In our model, the degree of local adaptation was spatially inconsistent. Argentinean phages (mainly from Chicoana) infected approximately as many local as nonlocal rhizobia, while Mexican phages were more infectious in local rhizobia than in nonlocal rhizobia. Spatial asymmetry in phage local adaptation is believed to be the result of the effects of nutrients on phage-host encounter rates, mutation rates and the cost of resistance [38, 101] or the local mode of coevolutionary dynamics (i.e., arms race or fluctuating selection [102]). Although we did not detect local phage maladaptation and we assumed that the differences in productivity across the sampled bean fields were insignificant, these studies show how environmental differences create spatially different intensities of phage local adaption. Indeed, the spatial heterogeneity of environmental factors results in a geographic mosaic of different evolutionary pressures [103]. Local adaptation to various environmental conditions can undermine the colonization success of allopatric individuals and limit gene flow (i.e., “isolation-by-adaptation,” or more general “isolation-by-environment” [33, 35, 101, 103]). Zhang & Buckling [34] found that host bacteria grown in the presence of phages in heterogeneous environments were limited in their ability to migrate across environments as a result of maladaptation. The limiting effect of local adaptation on phage migration has not been tested explicitly, although it has long been predicted [24, 101].

The spatial structure in the genetic composition of phage communities is probably due to the interplay of a variety of factors (*e.g*., historical contingencies, abiotic selection, genetic drift, and, potentially, dispersal limitation [8, 15]. Our results indicate that the presence of suitable hosts may play a role in shaping phage biogeography and that suitability is determined not only by the genetic identity of the host but also by local adaptation. The spatial patterns are analogous to those observed in *Rhizobium*-common bean interactions and suggest that the local adaptation of rhizobia to common bean may have shaped the spatial differences in the phage-rhizobium interactions. Through isolation-by-adaptation, local adaptation may reinforce spatial patterns in the phage community composition. Strong local adaptation of phages has been found across much shorter distances than in our present study [27, 96], and it is as yet unclear to what extent phage local adaptation leads to limited migration and at which scale this may occur. At smaller scales, spatial heterogeneity is probably under greater pressure due to higher viral migrant densities. However, across broad scales, local adaptation may be a significant barrier to successful long-distance phage migration.

## Acknowledgments

PAPIIT-UNAM IN209817 and the CCG-UNAM budget to VG funded the work. JVC received a Postdoctoral Scholarship from DGAPA-UNAM (2016-2018) and was partly supported by NSF grants DEB-1457508 and IOS-1759048, both awarded to E.L. Simms. We thank Gabriela Guerrero, José Espíritu, Alfredo Hernández, and Víctor del Moral for bioinformatics support, Alfonso Leija and Georgina Hernández for their help in greenhouse experiments, and Mario Marquina for providing access to the agricultural fields at Tepoztlán (Finca Xochitlamila) and Yautepec. We thank Ellen Simms, Olga María Pérez Carrascal, and Ellie Harrison for comments on previous versions of the manuscript. Special thanks go to Joshua Weitz for his advice on the BiMat application.

