## supplemental data for "Spatial patterns in phage-*Rhizobium* coevolutionary interactions across regions of common bean domestication"

Running title: Spatial patterns in phage-*Rhizobium* interactions.

Jannick Van Cauwenberghe [1,2 *], Rosa I. Santamaría [1, *], Patricia Bustos [1], Soledad Juárez [1], Maria Antonella Ducci[3], Trinidad Figueroa Fleming[4], Angela Virginia Etcheverry[4], and Víctor González [1].

[1] Centro de Ciencias Genómicas, Universidad Nacional Autonóma de México, Mexico**.**

[2] Department of Integrative Biology, University of California, Berkeley, CA, USA

[3] Instituto Nacional de Tecnología Agropecuaria, Universidad Nacional de Salta, Argentina.

[4] Facultad de Ciencias Naturales, Universidad Nacional de Salta, Salta, Argentina.

**Supplementary method S1**.

Obtention of *Rhizobium* strains from nodules.

Before inoculation with soil, bean seeds were surface sterilized in 70% ethanol for 1 minute and in commercial bleach (3% sodium hypochlorite) for 5 minutes, then finally rinsed six times with sterile water. Seeds were germinated on water agar plates at 30°C. Prior to inoculation, pots filled with 300 mL of air-dried vermiculite were autoclaved. Each pot was inoculated with a 100 mL soil sample, planted with 4 seedlings and received equal amounts of water. Each individual soil sample was divided into two pots: 3 populations * 3 soil samples/population * 4 plants/pot * 2 pots/soil sample = 72 plants. Plants received autoclaved water when necessary and were harvested after eight to twelve weeks of growth.

On average, three root nodules per individual plant were collected. The nodules were surface-sterilized in 70% ethanol for 1 minute and in commercial bleach (3% sodium hypochlorite) for 5 minutes, then finally rinsed six times with sterile water. Prior to the isolation of the rhizobia, root nodules were rolled over a peptone yeast extract agar plate containing nalidixic acid, $CaCl_2$ (7 mM) and $MgSO_4$ (10 mM) (*i.e.*, PYNal) to test the effectiveness of surface sterilization. The isolation of rhizobia was performed by squashing and streaking on PYNal agar plates. After an incubation period of three days at 30°C, all rhizobial isolates from successfully surface-sterilized nodules were purified by subculturing. The isolates were stored at -80°C in 50% glycerol for long-term storage.

Fig. S1. Map of the Americas showing the WGS 84 coordinates of the sampled common bean fields given in latitude and longitude (A) and the proportions of *Rhizobium* and phages isolated across the four common bean fields (B). The map was obtained from http://www.imagui.com/a/mapa-de-america-latina-sin-nombres-cdKbGxaoX.

**A**

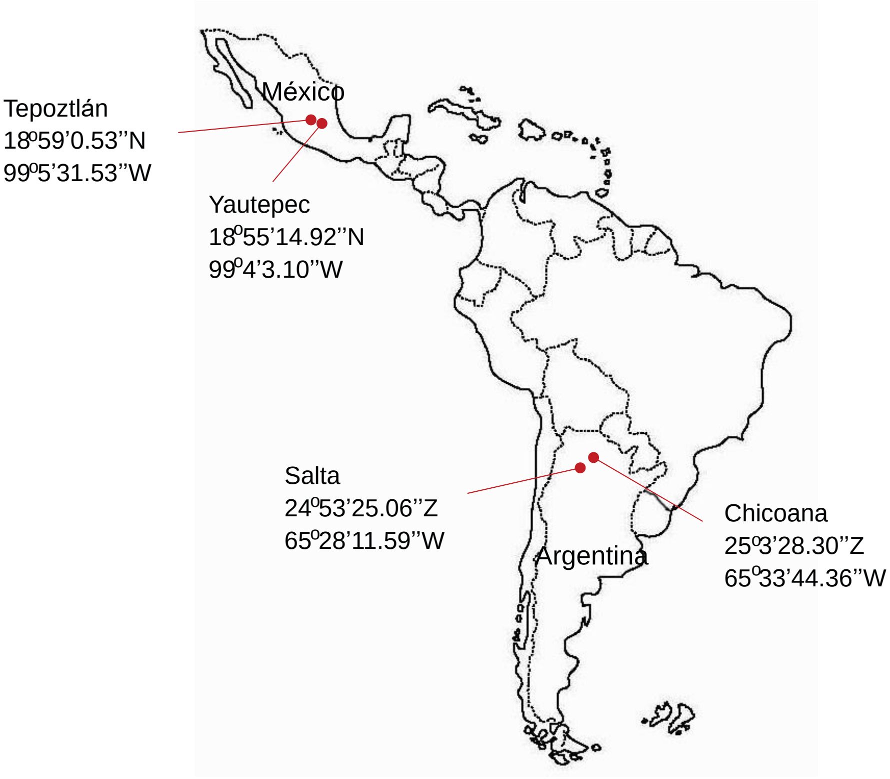


**B**

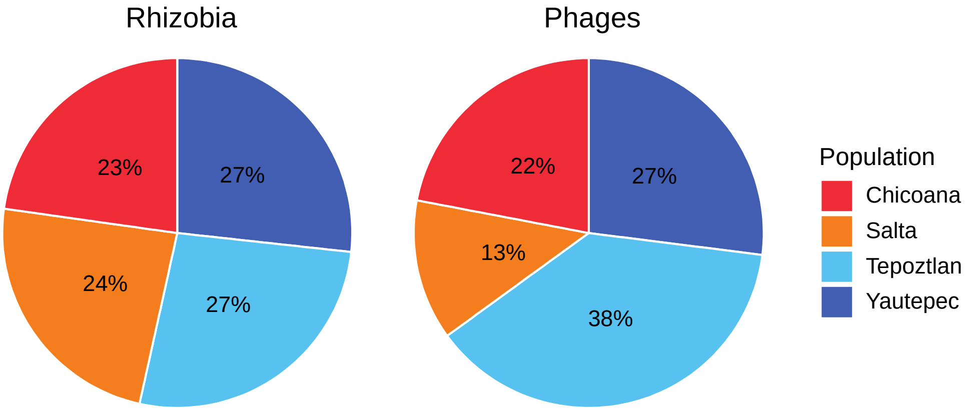

Fig. S2. Distribution of *Rhizobium* species (A) and phage families (B) across common bean fields.

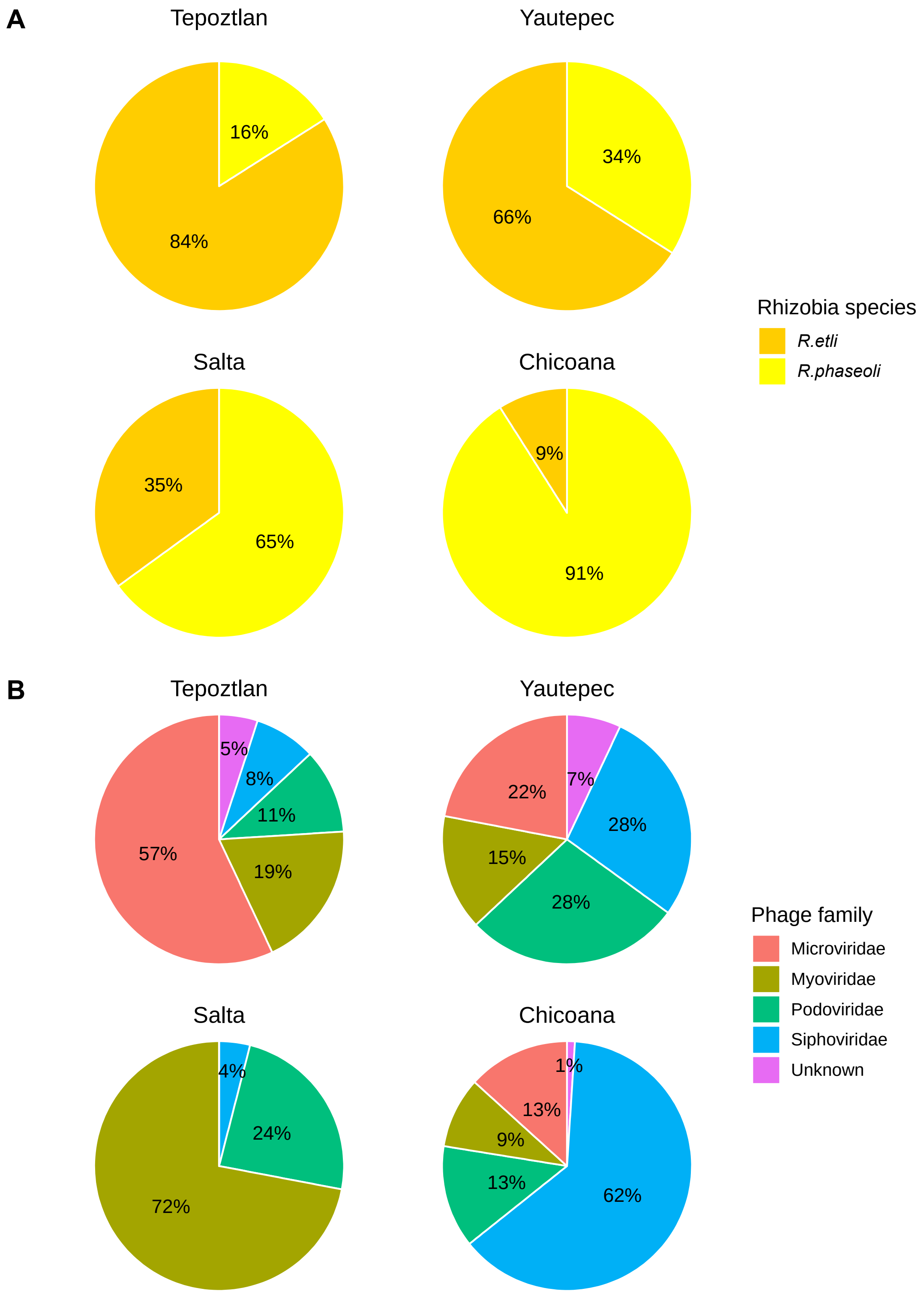

Fig S3. Heat map showing the products of ANIm (ANI by Mummer) and coverage. Phage families defined by ANIm and coverage. Genomes with a minimum of 80% nucleotide identity over an alignment covering at least 60% of the length of the smaller genome were grouped into a genomic family or phage genomic type (PGT; see Material and Methods).

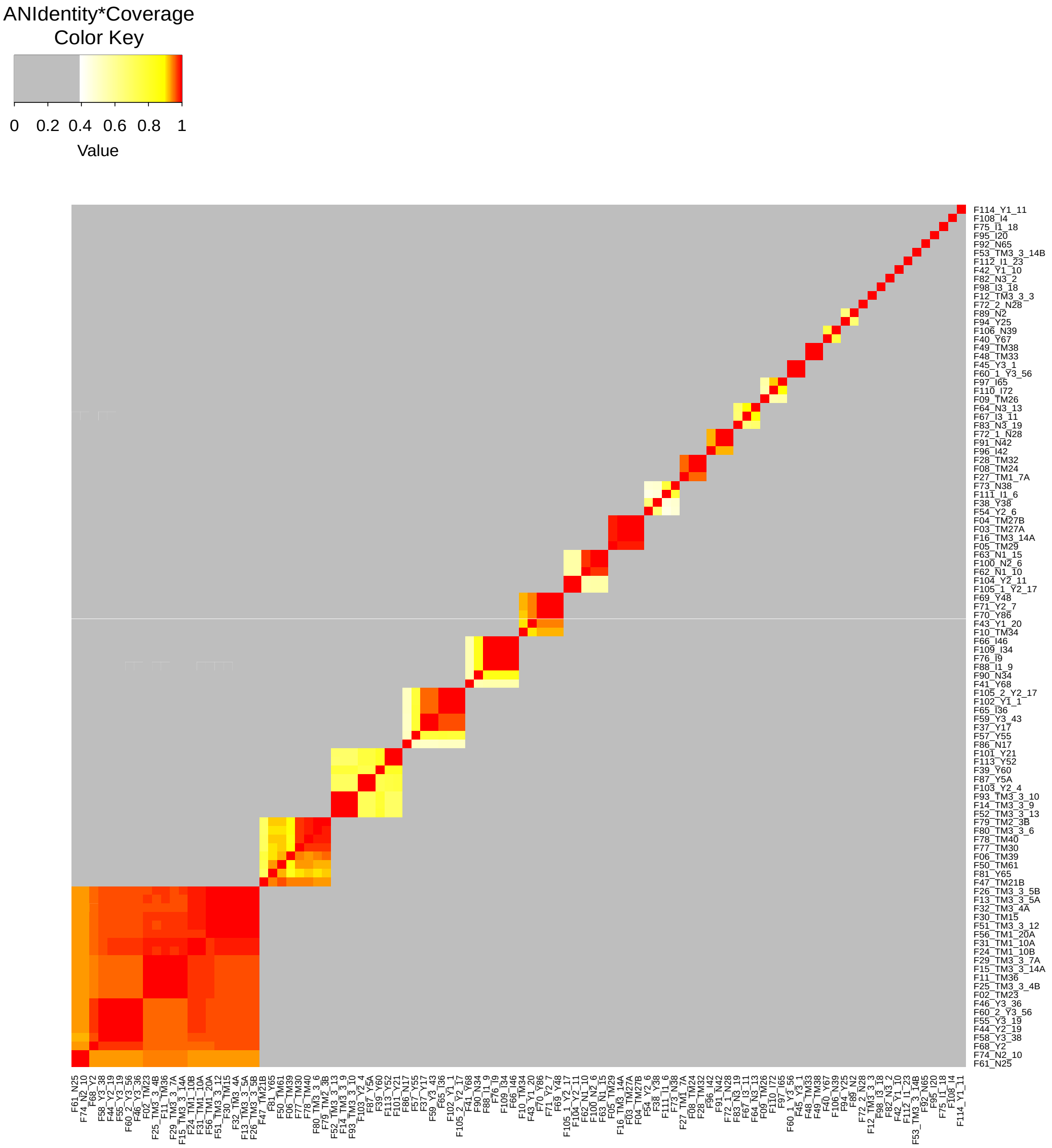

Table S1. List of SC strains with their taxonomic affiliation and the phages that were obtained using the respective strains.

| Code | Name | Species and symbiovar identity | Origin | # isolated phages |
|---|---|---|---|---|
| 1 | N1314 | *R. etli* sv. phaseoli | Tepoztlan, Morelos, Mexico | 2 |
| 2 | N741 | *R. etli* sv. phaseoli | Tepoztlan, Morelos, Mexico | 2 |
| 3 | N561 | *R. etli* sv. phaseoli | Tepoztlan, Morelos, Mexico | 2 |
| 4 | N771 | *R. phaseoli* sv. phaseoli | Tepoztlan, Morelos, Mexico | 2 |
| 5 | N161 | *R. phaseoli* sv. phaseoli | Tepoztlan, Morelos, Mexico | 5 |
| 6 | R72 | *R. gallicum* sv. phaseoli | Tepoztlan, Morelos, Mexico | 0 |
| 7 | R634 | *R. leguminosarum* sv. phaseoli | Tepoztlan, Morelos, Mexico | 1 |
| 8 | R744 | *R. phaseoli* sv. phaseoli | Tepoztlan, Morelos, Mexico | 1 |
| 9 | R635 | *R. leguminosarum* sv. phaseoli | Tepoztlan, Morelos, Mexico | 2 |
| 10 | N671 | *R. phaseoli* sv. phaseoli | Tepoztlan, Morelos, Mexico | 1 |
| 11 | N261 | *R. phaseoli* sv. phaseoli | Tepoztlan, Morelos, Mexico | 1 |
| 12 | R650 | *R. phaseoli* sv. phaseoli | Tepoztlan, Morelos, Mexico | 2 |
| 13 | R723 | *R. phaseoli* sv. phaseoli | Tepoztlan, Morelos, Mexico | 1 |
| 14 | R611 | *R. phaseoli* sv. phaseoli | Tepoztlan, Morelos, Mexico | 1 |
| 15 | R620 | *R. phaseoli* sv. phaseoli | Tepoztlan, Morelos, Mexico | 2 |
| 16 | R693 | *R. gallicum* sv. phaseoli | Tepoztlan, Morelos, Mexico | 1 |
| 17 | R630 | *R. phaseoli* sv. phaseoli | Tepoztlan, Morelos, Mexico | 4 |
| 18 | R711 | *R. gallicum* sv. phaseoli | Tepoztlan, Morelos, Mexico | 0 |
| 19 | N122 | *R. leguminosarum* sv. phaseoli | Tepoztlan, Morelos, Mexico | 0 |
| 20 | N324 | *R. leguminosarum* sv. phaseoli | Tepoztlan, Morelos, Mexico | 1 |
| 21 | N841 | *R. phaseoli* sv. phaseoli | Tepoztlan, Morelos, Mexico | 1 |
| 22 | N541 | *R. leguminosarum* sv. phaseoli | Tepoztlan, Morelos, Mexico | 1 |
| 23 | N621 | *R. etli* sv. phaseoli | Tepoztlan, Morelos, Mexico | 2 |
| 24 | N113 | *R. etli* sv. phaseoli | Tepoztlan, Morelos, Mexico | 1 |
| 25 | N6212 | *R. etli* sv. phaseoli | Tepoztlan, Morelos, Mexico | 4 |
| 26 | N4311 | *R. leguminosarum* sv. phaseoli | Tepoztlan, Morelos, Mexico | 1 |
| 27 | N831 | *R. phaseoli* sv. phaseoli | Tepoztlan, Morelos, Mexico | 3 |
| 28 | N731 | *R. etli* sv. phaseoli | Tepoztlan, Morelos, Mexico | 1 |
| 29 | N931 | *R. phaseoli* sv. phaseoli | Tepoztlan, Morelos, Mexico | 1 |
| 30 | N941 | *R. leguminosarum* sv. phaseoli | Tepoztlan, Morelos, Mexico | 2 |
| 31 | N1341 | *R. etli* sv. phaseoli | Tepoztlan, Morelos, Mexico | 0 |
| 32 | N871 | *R. etli* sv. phaseoli | Tepoztlan, Morelos, Mexico | 2 |
| 33 | R339 | *R. leguminosarum* sv. phaseoli | Tepoztlan, Morelos, Mexico | 2 |
| 34 | CFN42 | *R. etli* sv. phaseoli | Guanajuato, Mexico | 3 |
| 35 | CIAT652 | *R. phaseoli* sv. phaseoli | Costa Rica | 1 |
| 36 | GR56 | *R. phaseoli* sv. phaseoli | Spain | 2 |
| 37 | KIM5 | *R. phaseoli* sv. phaseoli | Idaho, USA | 2 |
| 38 | Bra5 | *R. phaseoli* sv. phaseoli | Brazil | 4 |
| 39 | S20 | *R. phaseoli* sv. phaseoli | Spain | 2 |
| 40 | CIAT894 | *R. leguminosarum* sv. phaseoli | Colombia | 2 |
| 41 | Mim1 | *R. etli* sv. mimosae | Huautla Morelos, Mexico | 1 |
| 42 | IE4771 | *R. phaseoli* sv. giardinii | Puebla, Mexico | 2 |
| 43 | IE4803 | *R. phaseoli* sv. phaseoli | Puebla, Mexico | 1 |
| 44 | IE4872 | *R. gallicum* sv. gallicum | Puebla, Mexico | 1 |
| 45 | IE951 | *R. phaseoli* sv. phaseoli | Puebla, Mexico | 3 |
| 46 | IE1004 | *R. phaseoli* sv. phaseoli | Puebla, Mexico | 3 |
| 47 | IE1006 | *R. phaseoli* sv. phaseoli | Puebla, Mexico | 0 |
| 48 | IE2737 | *R. phaseoli* sv. phaseoli | Puebla, Mexico | 1 |
| 49 | IE4777 | *R. phaseoli* sv. phaseoli | Puebla, Mexico | 0 |
| 50 | IE4810 | *R. phaseoli* sv. phaseoli | Puebla, Mexico | 0 |
| 51 | IE6766 | *R. gallicum* sv. phaseoli | Michoacan, Mexico | 0 |
| 52 | IE6794 | *R. leguminosarum* sv. phaseoli | Michoacan, Mexico | 1 |
| 53 | IE6833 | *R. etli* sv. phaseoli | Guanajuato, Mexico | 0 |
| 54 | IE6840 | *R. etli* sv. phaseoli | Guanajuato, Mexico | 0 |
| 55 | IE6845 | *R. etli* sv. phaseoli | Guanajuato, Mexico | 1 |
| 56 | IE6854 | *R. etli* sv. phaseoli | Guanajuato, Mexico | 0 |
| 57 | IE6868 | *R. etli* sv. phaseoli | Guanajuato, Mexico | 0 |
| 58 | IE6896 | *R. phaseoli* sv. phaseoli | Puebla, Mexico | 0 |
| 59 | CNPAF512 | *R. phaseoli* sv. phaseoli | Brazil | 0 |
| 60 | TAL182 | *R. etli* sv. phaseoli | Hawaii, USA | 1 |
| 61 | NXC12 | *R. etli* sv. mimosae | Huautla, Morelos, Mexico | 1 |
| 62 | NXC14 | *R. etli* sv. mimosae | Huautla, Morelos, Mexico | 0 |
| 63 | MIM2 | *R. phaseoli* sv. phaseoli | Huautla, Morelos, Mexico | 0 |
| 64 | MIM7-4 | *R. etli* sv. mimosae | Huautla, Morelos, Mexico | 0 |
| 65 | INC 1-2 | *R. phaseoli* sv. phaseoli | Guanajuato, Mexico | 4 |
| 66 | INC 2-5 | *R. phaseoli* sv. phaseoli | Guanajuato, Mexico | 1 |
| 67 | TUX10P | *R. phaseoli* sv. phaseoli | Tuxtlas Veracruz, Mexico | 1 |
| 68 | TUX16P | *R. leguminosarum* sv. phaseoli | Tuxtlas Veracruz, Mexico | 1 |
| 69 | TUX1712m | *R. phaseoli* sv. phaseoli | Tuxtlas Veracruz, Mexico | 0 |
| 70 | TUX250P | *R. phaseoli* sv. phaseoli | Tuxtlas Veracruz, Mexico | 0 |
| 71 | CH24-10 | *R. phaseoli* sv. phaseoli | Puebla, Mexico | 0 |
| 72 | 343 | *R. etli* sv. mimosae | | 1 |
| 73 | F6 | *R. etli* sv. phaseoli | | 0 |
| 74 | B27 | *R. phaseoli* sv. phaseoli | | 1 |
| 75 | BAT-4 | *R. etli* sv. phaseoli | Zacatecas, Mexico | 0 |
| 76 | BAT-49 | *R. phaseoli* sv. phaseoli | Zacatecas, Mexico | 0 |
| 77 | BAT-32 | *R. etli* sv. phaseoli | Zacatecas, Mexico | 0 |
| 78 | BAT-95 | *R. phaseoli* sv. phaseoli | Zacatecas, Mexico | 0 |
| 79 | D5 | *R. phaseoli* sv. phaseoli | Durango, Mexico | 0 |
| 80 | D11 | *R. phaseoli* sv. phaseoli | Durango, Mexico | 0 |
| 81 | D13 | *R. etli* sv. giardinii | Durango, Mexico | 0 |
| 82 | D21 | *R. etli* sv. giardinii | Durango, Mexico | 0 |
| 83 | PS14 | *R. phaseoli* sv. phaseoli | | 0 |
| 84 | 6C-1 | *R. gallicum* sv. phaseoli | Spain | 0 |
| 85 | 14C-1 | *R. phaseoli* sv. phaseoli | Spain | 0 |
| 86 | 14C-2 | *R. phaseoli* sv. phaseoli | Spain | 1 |
| 87 | 8NJ-2 | *R. etli* sv. phaseoli | Spain | 0 |
| 88 | 17NJ-2 | *R. gallicum* sv. phaseoli | Spain | 0 |
| 89 | 21NJ-2 | *R. gallicum* sv. phaseoli | Spain | 0 |
| 90 | 6PR-1 | *R. phaseoli* sv. phaseoli | Spain | 1 |
| 91 | GR10 | *R. gallicum* sv. phaseoli | Spain | 0 |
| 92 | GR14 | *R. phaseoli* sv. phaseoli | Spain | 0 |
| 93 | GR62 | *R. phaseoli* sv. phaseoli | Spain | 0 |
| 94 | GR75 | *R. phaseoli* sv. phaseoli | Spain | 0 |

Table S2. Primer sequences used for the partial amplification of the indicated *Rhizobium* and phage genes, for the taxonomic identification of rhizobia, and for the identification of phage genomic types. The following thermal cycling protocol was used: 94°C for 10 min, 30 X (94°C for 1 min, $T_a$ for 1 min and 72°C for 1 min) and 72°C for 10 min. $T_a$ is the annealing temperature: 57°C for rhizobium primers and 60°C for phage primers.

| ***Rhizobium sp.*** | **Gene target/annotation** | **Direction** | **Sequence (5´-3´)** |
|---|---|---|---|
| *Rhizobium sp.* | *nodC* | Forward | GCTTTCAAGGGCAGTAGCAG |
| | | Reverse | CCTCGGGAGTTCTGAAGATC |
| *Rhizobium sp.* | *dnaB* | Forward | CGCCGCTCGAAAGATTGCAG |
| | | Reverse | TCATCATCGAGCAGGCGAC |
| *Rhizobium sp.* | *recA* | Forward | GACAAAAGCAAGGCACTTGAA |
| | | Reverse | ACATCACGCCGATCTTCATGC |
| **Phage Genomic Type** | | | |
| F02 | Hypothetical | Forward | CGTTCACGAAAACRCGCACY |
| | | Reverse | TCGCATAGAACTKGTGGCGT |
| F03 | Putative coat protein | Forward | CGATATCAACCTCGGCGTA |
| | | Reverse | TTGTCGATGACCTTCCAGCC |
| F06 | Phage tail sheath protein | Forward | AGGTGGTGCTGATGGTTCTG |
| | | Reverse | TGGTGAATCTGCGACGAACA |
| F08 | Putative major capsid protein | Forward | GCCGGTTCATGGCGTATTTC |
| | | Reverse | GTGGCGTTCCAGTTGTTGAC |
| F10 | Putative major capsid protein | Forward | CGATACCATCATCCTGGCCC |
| | | Reverse | GATAGCGGCCTTGGGGATAC |
| F14 | Terminase | Forward | GGCGAACGTGAACAAGTTCC |
| | | Reverse | TGTTGAACCCGTCCAGATGG |
| F37 | Putative terminase protein large subunit | Forward | CAAGATYAACCTGCTYGCGC |
| | | Reverse | GGTTTGAGCGCAATGATCCC |
| F38 | Terminase subunit A | Forward | GAGCTTCCCGATACTTCCGG |
| | | Reverse | AGCTGGACACGAGCAGTAAC |
| F40 | Putative major capsid protein | Forward | TCTTCATCGATCCGGATGCG |
| | | Reverse | GAAACCCGGCTCCTTGAAGA |
| F41 | Tail completion and sheath stabilizer protein | Forward | GTCCACGAAACCATCTAACAACG |
| | | Reverse | GACGTCGACTTCCACGTTCT |
| F45 | putative hydrolase | Forward | TATCGGGATCCCCTACGTCC |
| | | Reverse | GGAGCCGATGAACACTCCAA |
| F62 | Putative tail assembly protein | Forward | CGACGTCGACTTCTACGAGG |
| | | Reverse | GATCATGCCCGGTTCGATCT |
| F64 | Major capsid protein | Forward | CAGTTCCGCAAGACCTCTGT |
| | | Reverse | CCAGTTGACATGGGACTGCT |
| F75 | DNA polymerase | Forward | TCATCGATCTCACACGCTCG |
| | | Reverse | GGTCTTGCGGCTTGTTGAAG |

Table S3. Groups of the phage host range defined as phage phenotype groups (PPGs), and groups of *Rhizobium* susceptibility expressed as *Rhizobium* phenotype groups (RPGs). The minimum and maximum number of infections, phage genomic types (PGT) per PPG, and *Rhizobium* STs per RPG are given.

| Phage Phenotypic Groups (PPGs) | | | | | *Rhizobium* Phenotypic Groups (RPGs) | | | | |
|---|---|---|---|---|---|---|---|---|---|
| | Phages | # min infections | # max infections | PGT´s (F´s) | | Strains | # min infections | # max infections | ST´s |
| PPG-1 | 42 | 97 | 132 | 2 & 8 | RPG-1 | 40 | 42 | 55 | 3, 5 & 41 |
| PPG-2 | 24 | 175 | 214 | 62 & 108 | RPG-2 | 38 | 105 | 142 | 5, 7, 10, 20, 26, 36, 39 |
| PPG-3 | 9 | 173 | 210 | 6, 10, 14 & 62 | RPG-3 | 36 | 87 | 145 | 21, 22, 33 & 34 |
| PPG-4 | 7 | 71 | 94 | 41 | RPG-4 | 19 | 48 | 75 | 14, 15, 16 & 17 |
| PPG-5 | 6 | 50 | 69 | 2, 14, 37 & 114 | RPG-5 | 12 | 56 | 63 | 18 |
| PPG-6 | 5 | 90 | 113 | 38 | RPG-6 | 9 | 93 | 137 | 5, 11, 27, 30 & 36 |
| PPG-7 | 4 | 73 | 90 | 3 | RPG-7 | 7 | 68 | 88 | 22, 27 & 34 |
| PPG-8 | 4 | 52 | 58 | 14 | RPG-8 | 5 | 84 | 122 | 22 & 32 |
| PPG-9 | 4 | 63 | 87 | 2 | RPG-9 | 5 | 58 | 68 | 23 & 35 |
| PPG-10 | 4 | 89 | 124 | 6, 37 & 62 | RPG-10 | 4 | 36 | 60 | 22, 33 & 34 |
| PPG-11 | 3 | 120 | 121 | 2 & 9 | RPG-11 | 4 | 50 | 69 | 13, 15 & 40 |
| PPG-12 | 3 | 107 | 116 | 38 & 40 | RPG-12 | 4 | 64 | 86 | 22 & 34 |
| PPG-13 | 3 | 41 | 42 | 45 | RPG-13 | 4 | 62 | 96 | 6 & 9 |
| PPG-14 | 3 | 136 | 151 | 6 | RPG-14 | 3 | 62 | 64 | 25 |
| PPG-15 | 3 | 76 | 88 | 2 | RPG-15 | 2 | 82 | 85 | 31 |
| PPG-16 | 2 | 151 | 152 | 62 | RPG-16 | 2 | 94 | 101 | 39 |
| PPG-17 | 2 | 51 | 56 | 41 | RPG-17 | 2 | 104 | 114 | 8 & 28 |
| PPG-18 | 2 | 83 | 87 | 2 | RPG-18 | 2 | 80 | 83 | 37 |
| PPG-19 | 2 | 9 | 12 | 2 | RPG-19 | 2 | 82 | 101 | 5 & 10 |
| PPG-20 | 2 | 113 | 120 | 10 | Singleton RPGs | 29 | 20 | 129 | 1, 2, 4, 5, 10, 11, 12, 19, 22, 23, 24, 29, 33, 34, 35, 36, 37, 38 |
| PPG-21 | 2 | 62 | 66 | 14 | | | | | |
| PPG-22 | 2 | 47 | 53 | 14 | | | | | |
| PPG-23 | 2 | 67 | 67 | 64 | | | | | |
| PPG-24 | 2 | 18 | 25 | 75 | | | | | |
| Singleton PPGs | 57 | 5 | 163 | 2, 3, 6, 9, 10, 12, 14, 37, 38, 40, 41, 42, 48, 53, 62, 64, 72_1, 72_2, 75, 82, 89, 92, 95, 98, 112 | | | | | |

Table S4. Relative abundance of chromosomal sequence types (ST) among common bean fields and regions.

| | | México | | Argentina | | | |
|---|---|---|---|---|---|---|---|
| ST code | Species | Tepoztlán | Yautepec | Salta | Chicoana | Mexico | Argentina |
| | | n=61 | n=61 | n=53 | n=54 | n=122 | n=107 |
| 1 | *R. phaseoli* | 0 | 0 | 2 | 0 | 0 | 1 |
| 2 | *R. phaseoli* | 0 | 0 | 0 | 2 | 0 | 1 |
| 3 | *R. phaseoli* | 2 | 0 | 0 | 0 | 1 | 0 |
| 4 | *R. phaseoli* | 2 | 0 | 0 | 0 | 1 | 0 |
| 5 | *R. phaseoli* | 2 | 30 | 19 | 83 | 16 | 50 |
| 6 | *R. phaseoli* | 0 | 0 | 2 | 0 | 0 | 1 |
| 7 | *R. phaseoli* | 0 | 0 | 2 | 0 | 0 | 1 |
| 8 | *R. etli* | 0 | 0 | 2 | 0 | 0 | 1 |
| 9 | *R. phaseoli* | 0 | 0 | 0 | 6 | 0 | 3 |
| 10 | *R. phaseoli* | 8 | 2 | 0 | 0 | 5 | 0 |
| 11 | *R. phaseoli* | 2 | 3 | 0 | 0 | 2 | 0 |
| 12 | *R. phaseoli* | 2 | 0 | 0 | 0 | 1 | 0 |
| 13 | *R. phaseoli* | 0 | 0 | 4 | 0 | 0 | 2 |
| 14 | *R. phaseoli* | 0 | 0 | 2 | 0 | 0 | 1 |
| 15 | *R. phaseoli* | 0 | 0 | 4 | 0 | 0 | 2 |
| 16 | *R. phaseoli* | 0 | 0 | 30 | 0 | 0 | 15 |
| 17 | *R. phaseoli* | 0 | 0 | 2 | 0 | 0 | 1 |
| 18 | *R. etli* | 0 | 0 | 22 | 0 | 0 | 11 |
| 19 | *R. etli* | 0 | 0 | 2 | 0 | 0 | 1 |
| 20 | *R. etli* | 0 | 0 | 0 | 2 | 0 | 1 |
| 21 | *R. etli* | 2 | 0 | 0 | 0 | 1 | 0 |
| 22 | *R. etli* | 23 | 8 | 0 | 0 | 16 | 0 |
| 23 | *R. etli* | 0 | 7 | 0 | 0 | 3 | 0 |
| 24 | *R. etli* | 2 | 2 | 0 | 0 | 2 | 0 |
| 25 | *R. etli* | 0 | 0 | 0 | 6 | 0 | 3 |
| 26 | *R. etli* | 0 | 2 | 0 | 0 | 1 | 0 |
| 27 | *R. etli* | 0 | 3 | 0 | 0 | 2 | 0 |
| 28 | *R. etli* | 0 | 0 | 2 | 0 | 0 | 1 |
| 29 | *R. etli* | 0 | 2 | 0 | 0 | 1 | 0 |
| 30 | *R. etli* | 0 | 5 | 0 | 0 | 2 | 0 |
| 31 | *R. etli* | 0 | 3 | 0 | 0 | 2 | 0 |
| 32 | *R. etli* | 7 | 0 | 0 | 0 | 3 | 0 |
| 33 | *R. etli* | 5 | 2 | 0 | 0 | 3 | 0 |
| 34 | *R. etli* | 46 | 11 | 0 | 0 | 29 | 0 |
| 35 | *R. etli* | 0 | 8 | 0 | 0 | 4 | 0 |
| 36 | *R. etli* | 0 | 7 | 0 | 0 | 3 | 0 |
| 37 | *R. etli* | 0 | 5 | 0 | 0 | 2 | 0 |
| 38 | *R. etli* | 0 | 2 | 0 | 0 | 1 | 0 |
| 39 | *R. etli* | 0 | 0 | 6 | 0 | 0 | 3 |
| 40 | *R. etli* | 0 | 0 | 2 | 0 | 0 | 1 |
| 41 | *R. etli* | 0 | 0 | 0 | 2 | 0 | 1 |

Table S5. Degrees of freedom (df), pseudo-F-values and P-values of PERMANOVA analyses testing the significance of compositional differences of *Rhizobium* chromosomal STs and phage PGTs across all four common bean fields and both common bean gene pools based on four distance matrices: Bray-Curtis dissimilarity matrix and Jaccard dissimilarity matrix. P-values for fitting the PCNM values on the PCoA are also given. Significant (P < 0.05) values are indicated in bold.

| Dataset | Distance matrix | Among fields | | | Among gene pools | | | PCNM fit |
|---|---|---|---|---|---|---|---|---|
| | | *df* | Pseudo -F- value | P-value | *df* | Pseudo -F- value | P-value | P-value |
| *Rhizobium* chromosomal STs | Bray-Curtis | 3,8 | 4.974 | **0.001** | 1,10 | 4.964 | **0.005** | **0.002** |
| | Jaccard | 3,8 | 3.561 | **0.001** | 1,10 | 3.363 | **0.005** | **0.001** |
| *R. phaseoli* chromosomal STs | Bray-Curtis | 3,8 | 2.385 | **0.009** | 1,10 | 1.969 | 0.070 | 0.116 |
| | Jaccard | 3,8 | 2.189 | **0.013** | 1,10 | 1.781 | 0.063 | **0.021** |
| *R. etli* chromosomal STs | Bray-Curtis | 3,6 | 4.081 | **0.009** | 1,8 | 3.959 | **0.001** | **0.002** |
| | Jaccard | 3,6 | 2.877 | **0.006** | 1,8 | 2.894 | **0.002** | **0.003** |
| Phage genomic types | Bray-Curtis | 3,10 | 3.265 | **0.001** | 1,12 | 3.379 | **0.001** | **0.001** |
| | Jaccard | 3,10 | 2.494 | **0.001** | 1,12 | 2.682 | **0.001** | **0.030** |

Table S6. Relative abundance of phage genomic types among common bean fields and regions. Taxonomic family identities are given.

| | | México | | Argentina | | México | Argentina |
|---|---|---|---|---|---|---|---|
| **PGT** | **Taxonomic Family** | **Tepoztlán n= 74** | **Yautepec n= 50** | **Salta n= 45** | **Chicoana n= 26** | **n= 124** | **n=71** |
| F02 | Microviridae | 49 | 22 | 0 | 11 | 38 | 7 |
| F03 | Podoviridae | 7 | 0 | 0 | 0 | 4 | 0 |
| F06 | Myoviridae | 19 | 2 | 0 | 0 | 12 | 0 |
| F08 | Microviridae | 12 | 0 | 0 | 0 | 7 | 0 |
| F09 | Myoviridae | 1 | 0 | 8 | 0 | 1 | 3 |
| F10 | Podoviridae | 1 | 12 | 0 | 0 | 6 | 0 |
| F12 | Siphoviridae | 1 | 0 | 0 | 0 | 1 | 0 |
| F14 | Siphoviridae | 5 | 16 | 0 | 0 | 10 | 0 |
| F37 | Myoviridae | 0 | 12 | 4 | 2 | 5 | 3 |
| F38 | Podoviridae | 0 | 12 | 4 | 2 | 5 | 3 |
| F40 | Microviridae | 0 | 2 | 0 | 2 | 1 | 1 |
| F41 | Myoviridae | 0 | 4 | 42 | 2 | 2 | 17 |
| F42 | Podoviridae | 0 | 2 | 0 | 0 | 1 | 0 |
| F45 | Siphoviridae | 0 | 8 | 0 | 0 | 3 | 0 |
| F48 | Podoviridae | 3 | 0 | 4 | 2 | 2 | 3 |
| F53 | Siphoviridae | 1 | 0 | 0 | 0 | 1 | 0 |
| F62 | Siphoviridae | 0 | 4 | 0 | 60 | 2 | 38 |
| F64 | Podoviridae | 0 | 0 | 8 | 4 | 0 | 6 |
| F72_1 | Myoviridae | 0 | 0 | 4 | 4 | 0 | 4 |
| F72_2 | Siphoviridae | 0 | 0 | 0 | 2 | 0 | 1 |
| F75 | Myoviridae | 0 | 0 | 12 | 0 | 0 | 4 |
| F82 | Podoviridae | 0 | 0 | 0 | 2 | 0 | 1 |
| F89 | Podoviridae | 0 | 2 | 0 | 2 | 1 | 1 |
| F92 | Podoviridae | 0 | 0 | 0 | 2 | 0 | 1 |
| F95 | Podoviridae | 0 | 0 | 4 | 0 | 0 | 1 |
| F98 | Podoviridae | 0 | 0 | 4 | 0 | 0 | 1 |
| F108 | Siphoviridae | 0 | 0 | 4 | 0 | 0 | 1 |
| F112 | Podoviridae | 0 | 0 | 4 | 0 | 0 | 1 |
| F114 | Siphoviridae | 0 | 2 | 0 | 0 | 1 | 0 |

Table S7. Mantel's r statistics and P-values for correlations among compositional differences of *Rhizobium* chromosomal STs, PGTs, RPGs, and PPGs across all four common bean fields. Correlations using the Jaccard distance matrix (above the diagonal, in gray), and the Bray-Curtis distance matrix (below the diagonal).

| | **Data set** | | | |
|---|---|---|---|---|
| **Data set** | **STs** | **PGTs** | **RPGs** | **PPGs** |
| **STs** | - | r= 0.540; P = 0.003 | r = 0.681; P = 0.001 | r = 0.682; P = 0.002 |
| **PGTs** | r= 0.433; P = 0.002 | - | r= 0.400; P = 0.014 | r= 0.628; P = 0.001 |
| **RPGs** | r = 0.870; P = 0.001 | r= 0.439 ; P = 0.009 | - | r = 0.523; P = 0.004 |
| **PPGs** | r = 0.457; P = 0.007 | r= 0.764; P = 0.001 | r = 0.467; P = 0.010 | - |
